# Modulation of quantitative trait loci for *Arabidopsis thaliana* seed performance by the maternal and germination environment

**DOI:** 10.1101/2023.02.22.529582

**Authors:** Basten L. Snoek, Elise A. R. Serin, Harm Nijveen, Leo A. J. Willems, Juriaan A. Rienstra, Martijn van Zanten, Henk W. M. Hilhorst, Wilco Ligterink

## Abstract

The quality of seeds contributes to plant performance, especially during germination and in the young seedling stage, and hence affects the economic value of seed crops. A seed’s innate quality is determined during seed development and the following seed maturation phase. It is tightly controlled by the genetic make-up of the mother plant and further shaped by the environmental conditions of the mother plant. The interaction between genotype and environment can result in substantial quantitative variation in seed traits like dormancy and viability.

Making use of naturally occurring variation within the *Arabidopsis thaliana* germplasm, we studied the interaction between seed production environments and the genetic architecture of mother plants on diverse seed quality traits. An Arabidopsis Bayreuth-0 x Shahdara recombinant inbred line (RIL) population was grown in four different seed production environments: high temperature, high light, low phosphate, and control conditions. The seeds harvested from the mother plants that were exposed to these environments from flowering until seed harvest were subsequently subjected to germination assays under standard and mild stress conditions (cold, heat, osmotic stress and added phytohormone ABA). Quantitative trait locus (QTL) analysis identified many environmental-sensitive QTLs (QTL x E) as well as several interactions between the maternal and germination environments. Variation in the number and position of the QTLs was largely determined by the germination conditions, however effects of the maternal environment were clearly present regarding the genomic location as well as significance of the individual QTLs.

Together, our findings uncover the extensive environmental modulation of the genetic influence on seed performance and how this is shaped by the genetic make-up of the mother plant. Our data provides a systems-view of the complex genetic basis of genotype-by-environment interactions determining seed quality.

## Introduction

Seed performance traits such as dormancy, viability and vigor are critical in the early stages of the plant’s life cycle as they ultimately determine the fitness of the individual and reproductive success within a population [1]. Seed performance is determined by the combination of environmental conditions and the quality of the seeds [2, 3]. The latter has a genetic basis and is established during seed development and maturation, occurring mostly on the mother plant before shedding [2, 3]. Important seed quality characteristics include: i) the ability of the seeds to germinate timely and uniformly under a wide range of conditions (vigor, seed dormancy), ii) genetic purity, iii) ability to be stored (in the soil or on the shelf) for a long period of time without losing viability (longevity) and iv) the capability to establish a strong and resilient seedling [4]. In agriculture, the quality of seeds scales to the performance of crops [5].

The timing of germination, an important aspect of seed quality, is controlled by innate seed dormancy. Seed dormancy is defined as the inability to germinate shortly after maturation, despite favorable environmental conditions for germination and allows the seed to overcome upcoming unfavorable periods - such as winter cold - for seedling establishment [6] [7]. Primary dormancy in Arabidopsis can be released by a period of dry storage termed after-ripening. In many natural accessions, a quick release of dormancy can be accomplished by a combined dark and cold treatment (stratification) applied to imbibed seeds prior to initiation of germination. Another important determinant of the success of germination is the post-dispersal environment of non-dormant seeds. Unfavorable germination conditions like dry, hot, or saline soils, can reduce, delay, or prevent seed germination [8].

Seed development and maturation are critical phases in a (mother) plant’s life cycle, yet are particularly sensitive to environmental perturbations, including stresses [9, 10]. Both at the vegetative and reproductive stage, the mother plant, as well as the developing seed, process environmental information [11]. Environmental factors such as temperature [12–15], light quality and intensity [2, 16, 17], photoperiod [18] and nutrient availability [2, 19] affect many plant and seed traits. Interestingly, several studies suggest that environmental conditions perceived by the mother plant affect the timing of germination of its offspring seeds [2, 20–22]. For instance, perception of high temperature during seed development can result in reduced seed dormancy [14], while seed maturation under cold conditions can induce strong dormancy [20, 23].

The occurrence of extensive natural phenotypic variation in diverse seed traits sparked investigations into the genetic basis of traits like dormancy [24, 25] and germination in Arabidopsis [8, 26–29] and in commercial crops [30, 31]. Genetic variation was also observed for the effect of the maternal environment on seed traits, in panels of different genotypes [18]. Significant genotype x environment (GxE) interactions were detected [2, 32–34], suggesting that the genetic contribution to GxE interaction can be unraveled by quantitative genetic approaches. This genetic contribution can be estimated by assessing recombinant inbred line (RIL) populations derived from genetic backgrounds that segregate for the traits of interest, followed by quantitative trait locus (QTL) mapping [35]. Testing RIL populations in multiple environments can subsequently bring substantial insight into the QTL/GxE interactions underlying the phenotypic expression of traits under study [8, 36–42].

The effects of the maternal environment on the genetic architecture of seed quality and germination traits are still sparsely studied as only a few publications provide insight into the GxE interaction of the maternal environment and how this shapes the QTL landscape of offspring seed quality [43–45]. The focus of these studies was mainly on seed dormancy, in view of the trait’s ecological implications [20]. Additional knowledge on diverse seed quality traits, including seed vigor, can provide a more comprehensive understanding of seed performance and its plasticity, potentially to be used in plant breeding efforts towards the development of more resilient varieties [5, 32, 33].

In this study, we used an *Arabidopsis thaliana* RIL population derived from two natural accessions: Bayreuth-0 (Bay-0) and Shahdara (Sha) [46], to probe the interactions between the maternal environment and the genetic architecture of the mother plants on seed quality and germination traits of seeds exposed to different environmental conditions. The parental lines and the RIL population were grown in four different seed production environments: high temperature, high light, low phosphate and standard (control) conditions, from flowering until seed harvest [2]. Germination characteristics of the harvested seeds were quantified following exposure to diverse environmental germination conditions (imbibed/non-imbibed, stratified/non-stratified, dry/imbibed, cold, heat and high salinity). This enabled us to describe the interactions between genetic background, maternal and germination environment. Our work uncovers a complex genetic architecture with both robust and highly environment-specific QTLs for seed and germination traits. The detection of extensive interaction between genotype, maternal and germination environment resulting in plasticity in seed development and germination can be of use in targeted breeding approaches aiming for resilient crop varieties that perform optimally when exposed to specific environmental perturbations.

## Results

### Phenotypic variation in seed quality and germination traits

To determine the genetic basis of diverse seed and germination traits of seeds produced under different maternal environments (ME), a Bay-0 x Sha recombinant inbred line (RIL) population [46, 47] was grown in standard (ST) long day conditions (light intensity of 150 μmol m s, day/night cycle of 16h/8h at 22°C/18°, 0.5 mM P). From the moment of floral initiation, ‘mother’ plants were moved to either of three controlled mild stress environments: high temperature (HT; day/night cycle of 25°C/23°C), high light (HL; 300 μmol m s) or low phosphate (LP; 0.01 mM P) or were kept in the previously mentioned standard conditions (ST) as a control, until the ripe seeds were harvested. Using the harvested seeds, we quantified 70 traits including seed dormancy, seed longevity and seed vigor as well as germination rates (**Supplemental Table 1-3**). For most seed and germination traits extensive phenotypic variation was observed within the RIL population and among the parental lines in the different environments (**Figure 1, Figure 2, Supplemental Figure 1, Supplemental Table 3**).

**Figure 1.**
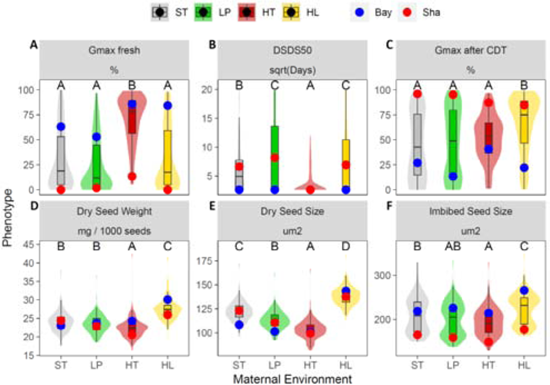
Effect of the seed maternal environment on offspring seed and germination traits of RIL population genotypes and parental lines. The phenotypic values of the parental lines, Bay-0 and Sha are indicated with blue and red dots, respectively. (A) percentage of germination of freshly harvested seeds (G_max_ fresh). (B) number of days of dry storage of seeds required to reach 50% of germination (DSDS_50_). DSDS_50_ data was square root transformed. (C) Germinating fraction (percentage) after controlled deterioration (CDT), a proxy for seed longevity (G_max_ after CDT). (D) Average dry seed weight of 1000 seeds in milligram (mg) (Dry seed weight). (E) Average projected seed size of 1000 seeds in micrometer^2^ (µm2) (Dry seed size). (F) Average projected size of imbibed seeds in micrometer^2^ (µm2) (Imbibed seed size). Boxes indicate boundaries of the second and third quartiles of the RIL distribution data. Black horizontal bars indicate median and whiskers Q1 and Q4 values within 1.5 times the interquartile range. Violin plots designate phenotype distributions. Colored shadings represent the different maternal environments; standard (control) conditions (ST, grey), low phosphorus (LP, green), high temperature (HT, red) and high light (HL, yellow). Significant differences between maternal environments are indicated by different capital letters above the plots, calculated using an ANOVA test with post-hoc Tukey HSD (p-value < 0.01).

**Figure 2.**
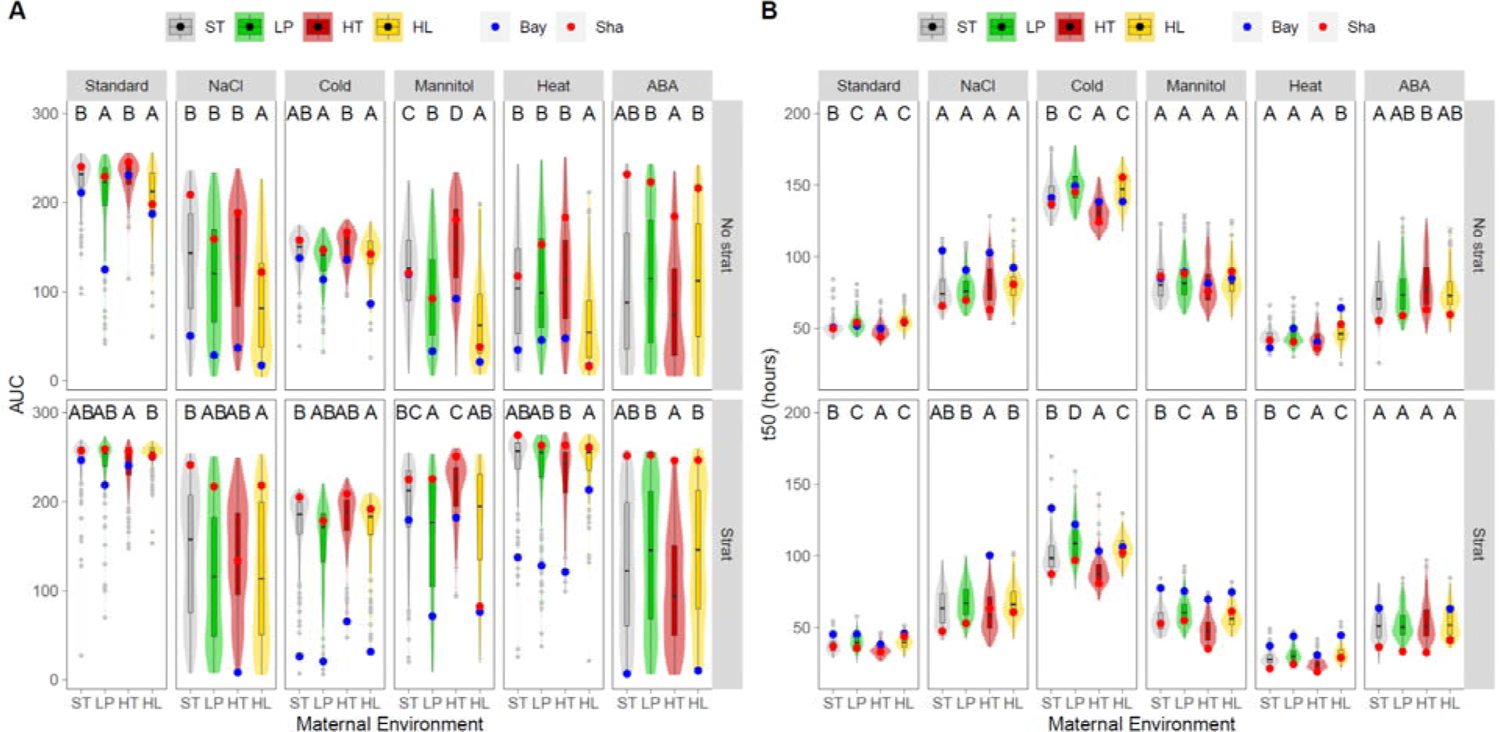
Effect of the seed maternal and germination environments on seed vigor traits of RIL population genotypes and parental lines. (A) AUC and (B) t_50_ values. The phenotypic values of the parental lines, Bay-0 and Sha are indicated with blue and red dots, respectively. AUC and t_50_ were assessed using fully after-ripened non-stratified seeds (upper row) and fully after-ripened stratified seeds (lower row) and were exposed to salt, cold, heat or osmotic stress or ABA. Boxes indicate boundaries of the second and third quartiles of the RIL distribution data. Black bars indicate median and whiskers Q1 and Q4 values within 1.5 times the interquartile range. Outliers are represented by grey dots. Violin plots designate phenotypic distributions. Colored shadings represent the different maternal environments; standard (control) conditions (ST, grey), low phosphorus (LP, green), high temperature (HT, red) and high light (HL, yellow). Significant differences between maternal environments are indicated by different capital letters above the plots calculated using an ANOVA test with post-hoc Tukey HSD (p-value < 0.01).

Compared to Bay-0, Sha seeds derived from mother plants of all maternal environments had a higher level of primary dormancy (G_max_ fresh; **Figure 1A**), indicated by a lower percentage of germination of fresh harvested seeds, and by its requirement of more days of seed dry storage to reach 50% germination (DSDS_50_; **Figure 1B**). Imbibed Sha seeds derived from all ME’s also displayed a higher fraction of germinated seeds after controlled deterioration treatment (G_max_ CDT; **Figure 1C**), indicative for a higher longevity. In standard conditions (ST), Sha seeds were slightly heavier than those of Bay-0, yet under all three mild stress ME’s Bay-0 seed weight superseded those of Sha (Dry seed weight; **Figure 1D**). This was partly reflected in larger seeds, with Sha seeds being bigger in the ST and LP environment, but not when derived from the HT or HL environment (Dry seed size; **Figure 1E**). However, after imbibition seeds of Bay-0 were larger regardless of the ME (Imbibed seed size; **Figure 1F**). Considering the whole RIL population, the HL maternal environment had the most pronounced effect on seed quality and germination traits as compared to ST control conditions. The HT maternal environment led to an increase in primary dormancy (G_max_ fresh), reduced DSDS_50_, and reduced seed size of both dry and imbibed offspring seeds (**Figure 1A, B, E, F**). The HL maternal environment led to an overall increased DSDS_50_, reduced longevity and increases in seed weight and size (**Figure 1B, C, E, F**). The LP maternal environment had relatively mild effects and only resulted in increased DSDS_50_ and mildly reduced seed size (**Figure 1B, F**).

For most traits some transgression (offspring with phenotypic values extending beyond the parental values) was observed **(Figure 1)**. For G_max_ transgression was one-sided, where a part of the RILs showed higher values than the highest-value parental line (Bay-0 in this case) but did not exceed the Sha value on the other side of the trait distribution (**Figure 1A)**. For the HT maternal environment this was less pronounced as the G_max_ was overall increased in this condition. One-sided transgression was also found for DSDS_50_ (**Figure 1B)**, for which also HT had a pronounced effect, lowering the DSDS_50_ for most RILs. Two-sided transgression was found for the traits G_max_ after CDT, dry seed weight, dry seed size, and imbibed seed size (**Figure 1C-F)**. For these traits the HL maternal environment had the strongest effect on the phenotypic distribution in the RILs. A smaller but noticeable effect was also found for the HT environment. Together this shows complex interactions between the genetic background, maternal environments, and specific traits (**Figure 1)**.

Seed vigor was assessed by assaying diverse seed germinating traits of fully after-ripened stratified and non-stratified seeds exposed to different germination environments (GE; **Supplemental Table 1, 2**). GE’s included were salt (100 mM NaCl), cold (10°C), heat (32°C) and osmotic stress (−0.6 MPa mannitol) and the presence of the phytohormone Abscisic acid (0.25 µM ABA). The latter is associated with suppression of germination and induction of seed dormancy [6]. Standard conditions (ST, in water at 20°C) were used as control. The effect of these GE’s was tested by measuring germination rate (G_max_), germination speed (t_10_ and t_50_; time needed to reach 10% or 50% of total germination), uniformity of germination (u_8416_; the time interval between 16% and 84% of viable seeds to germinate) and the area under the germination curve until 300 hours (AUC), summarizing the mentioned germination parameters (**Supplemental Table 1, 2, 3**).

As found for seed traits (**Figure 1**), extensive phenotypic variation was observed for the diverse germination traits among the genotypes and parental lines across ME’s and GE’s (**Figure 2, Supplemental Figure 1, Supplemental Table 2**). We first determined the (dis)similarity in variation between the diverse traits within the RIL population (and their parental lines) by comparing the patterns found by Principal Component Analysis (PCoA) (**Supplemental Figure 2**). This pointed to the existence of two major trait groups in our data. On one hand the t_10_, t_50_ and u_8416_ clustered together and on the other hand AUC and G_max_ (**Supplemental Figure 2**). We therefore focused on the distributions of t_50_ and AUC values, as representative proxy of the two trait groups. The data of the other traits (G_max_, u_8416_ and t_10)_ are provided in **Supplemental Figure 1** and all data (including AUC and t_50_) can be accessed and further explored interactively through the AraQTL website (www.bioinformatics.nl/AraQTL/; [48]).

Overall, AUC values of the Bay-0 parental line were lower than those of Sha in most GE’s across ME’s (**Figure 2A**) in both stratified and non-stratified seeds, indicating that Sha seeds are more vigorous and resilient against sub-optimal germination conditions. This is in line with the higher stress sensitivity reported for Bay-0 as compared to Sha [8, 29]. We observed that most of the RILs (> 80%) did germinate readily in standard conditions (ST; i.e, water at 20°C) with little variation, while germination in mild stress-inducing environments was overall reduced but exhibited larger phenotypic variation across the RIL population. Although seed vigor was affected by the maternal environment (ME) to different extent, the germination environment overall appears much more determinative for progeny seed vigor. Put in other words, the difference between ST and average trait effect of individual GE’s is larger than the trait variation observed between ME’s within a GE block. Under some conditions, marked transgression was observed, as substantial parts of the segregating progenies (RILs) were clearly performing worse or better than the parental genotypes (**Figure 2A**).

In most GE’s, Bay-0 showed higher t_50_ values than Sha, indicating that Sha seeds germinated quicker (**Figure 2B**). This is in line with previously reports [8, 29]. Considering the GE’s, overall slowest germination was observed under cold conditions and fastest germination under high temperatures. In all cases, the effect of the GE prevailed over the effect of the ME (**Figure 2B**). Nevertheless, several significant effects of the maternal environment were detected. Transgression was predominantly two-sided and was observed in all GE and ME combinations (**Figure 2B**). Of note, overall, the germination environment is more determinative for AUC as well as t_50_ values than the maternal environment for both stratified and non-stratified seeds.

To probe for signs of possible shared genetic architecture, correlations between the diverse seed and germination trait values obtained from the different GE’s and ME’s across the RILs and parental lines were calculated (**Figure 3, Supplemental Table 4**). In general, only moderate correlations between the diverse seed traits, such as size, and seed germination traits were observed, suggesting limited overlap in determinative genetic components. Of note, the trait correlations were overall stronger when traits were obtained from non-stratified seeds as compared to those of stratified seed batches, suggesting that stratification in part dampens the effect of the ME and/or GE. Seed dormancy (DSDS_50_) was as expected generally negatively correlated with AUC (**Figure 3A**) and positively with t_50_ (**Figure 3B**) and LP and HL maternal environments exacerbated this effect. On the contrary, G_max_ CDT demonstrated a positive interaction with AUC, and negative with t_50_, which was exacerbated under HT (**Figure 3**). No particularly clear correlation was observed between dry seed size and seed vigor (AUC and t_50_) traits.

**Figure 3:**
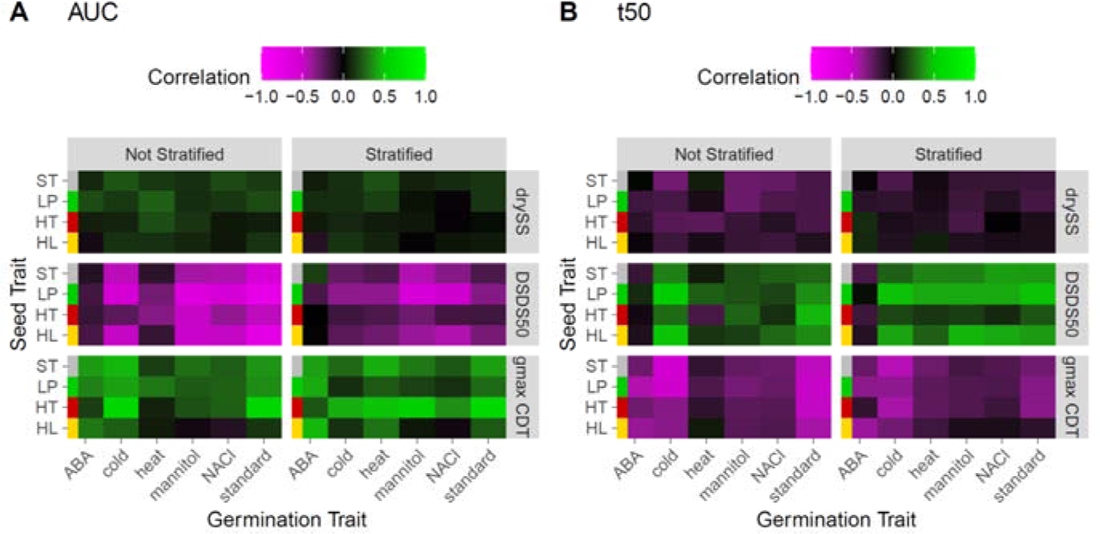
Heatmap of Spearman correlation coefficients of RIL seed and germination trait values. **(A)** AUC and **(B)** t_50_ correlation values obtained after exposure to different germination conditions. The x-axes (columns) represent different GE’s: Standard conditions, salt, cold, heat or osmotic stress or ABA application. The y-axis (rows) represents ME’s: standard, ST, grey color coding; low phosphate, LP, green; high temperature, HT, red; high light, HL, yellow. Seed traits included are number of days of dry storage of seeds required to reach 50% of germination (DSDS_50_), percentage of germination after controlled deterioration (G_max_ CDT) and dry seed size (drySS). Correlations for both stratified and non-stratified seeds are shown.

### G x E interactions for seed germination traits

As a result of genotype by environment (G x E) interactions, trait values per genotype (in our case individual RILs and parental lines) can differ from one environment to another. This can cause the genotypes to rank differently on phenotypic values between the environments, depending on the relative contribution of underlying genetic components vs. the effect of the environmental condition [33]. To estimate G x E interactions in our dataset, Spearman correlation analysis and clustering of germination traits across the multiple environments was performed for all ME x GE combinations and all RIL and parental genotypes (**Figure 4, Supplemental Table 4**). Overall, positive correlations of the AUC phenotype between the ME and GE conditions were found (**Figure 4A**). Also, for t_50_ mostly positive correlations between the ME’s and GE’s was observed (**Figure 4B**). This points to a common genetic basis, within the t_50_ traits and within the AUC traits, independent of the ME and GE, as well as a part that can be ME and/or GE specific (**Figure 4**). Principle Component Analysis confirmed the substantial environmental contribution to trait values (**Figure 5**). Variation in the AUC seed germination traits between different genetic backgrounds (RILs and parental lines) were mainly explained by the germination environment (e.g. ABA), but the maternal environment conditions also caused some recognizable clustering, mainly attributed to the HT maternal environment. This suggests additional effects of the maternal (ME) over the germination environment (GE) (**Figure 5**). For t_50_, a clear separation was identified for stratified vs. non-stratified seeds (**Figure 5B**), which was less-so the case for AUC (**Figure 5A**).

**Figure 4.**
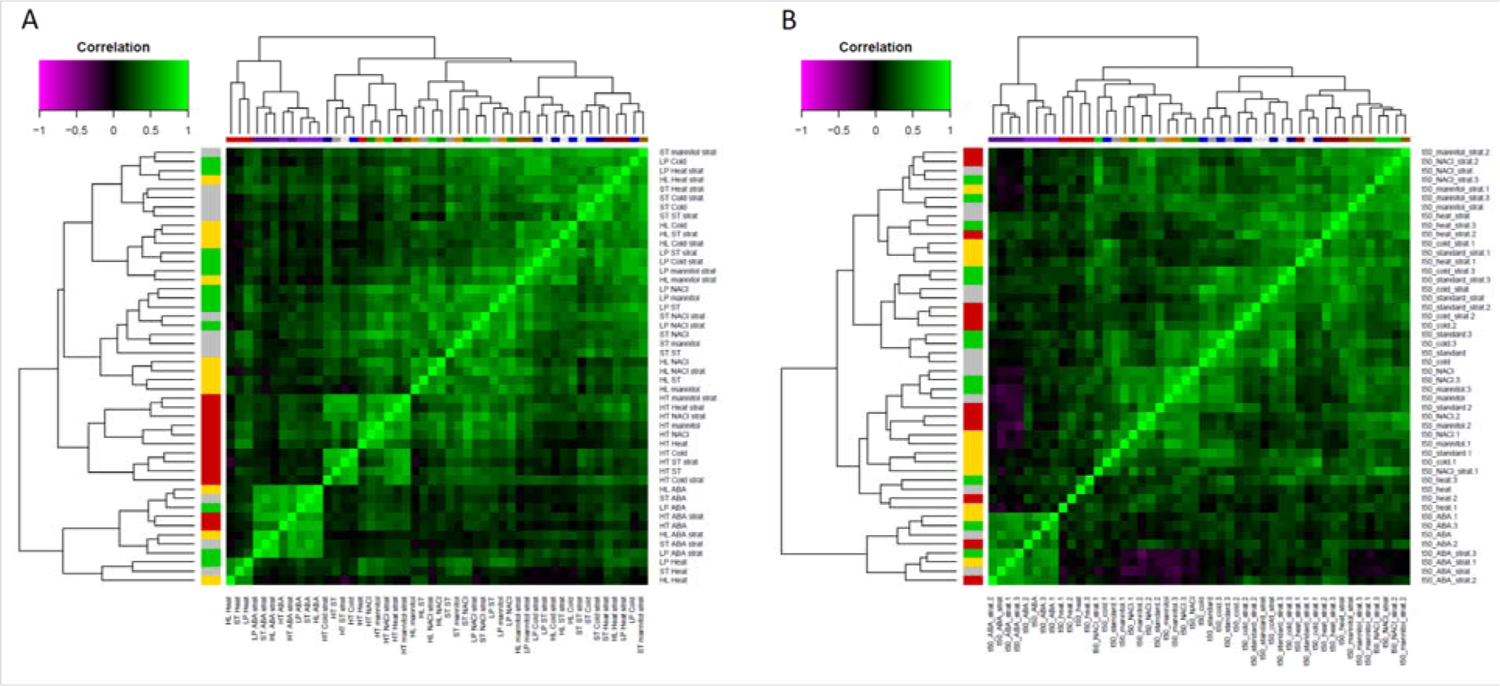
Heatmap and hierarchical clustering of S correlation coefficients of **(A)** AUC and **(B)** t_50_ seed germination trait values among all tested maternal and seed germination environments. Negative correlations are indicated in purple shading, positive correlations in green shading (see legend at the top left). The 48 possible maternal and seed germination environments are indicated on the axes and color coded. Maternal environments indicated on the y-axis are coded: ST, grey; LP, green; HT, red; HL, yellow. The germination environments on the top x-axis are coded: standard, (ST) grey; ABA, purple; cold, blue; heat, red; mannitol, yellow/brown; NaCl, green. Stratified seeds are indicated with dark colors and with ‘strat’. The remaining category of non-stratified seeds is not letter-coded but is indicated by lighter colors.

**Figure 5:**
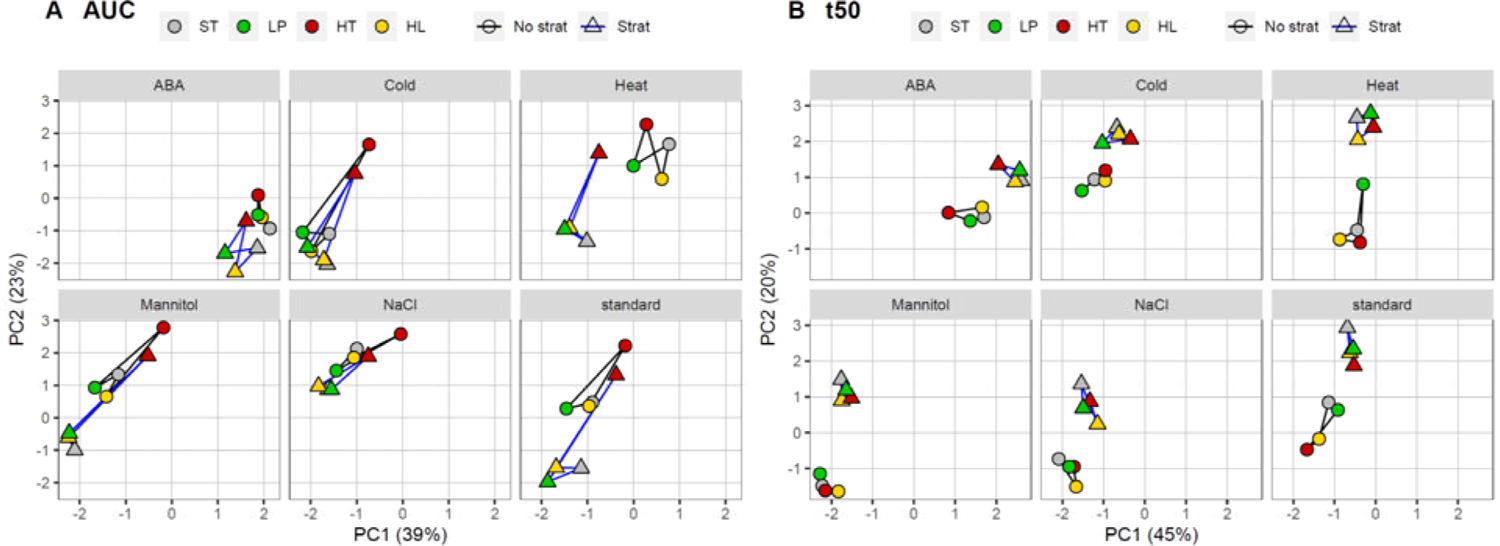
Biplots of the first two Principal Component (PC) axes of (**A**) AUC and (**B**) t_50_ trait value distributions within the RIL population, including the parental lines (all tested maternal environments and stratified and non-stratified seeds), plotted per germination environment (indicated above the panels). Maternal environment conditions are indicated; ST, grey; HL, green; HT, red; LP, yellow. Stratified seeds are indicated as triangles, non-stratified seeds as circles. Lines connect the samples per stratification treatment (grey, not stratified; blue; stratified).

### QTL mapping of seed traits

To define the genetic architecture of the contribution of the maternal environment (ME) to diverse seed traits, we adopted an QTL mapping approach. Mapped traits include imbibed and dry seed size (SS), dry seed weight (SW), percentage of germination of fresh seeds (G_max_) and after controlled deterioration (G_max_ CDT), and days of dry storage required to reach 50% of germination (DSDS_50_). The obtained QTL profiles per ME were overlayed to find differences and similarities (**Figure 6, Supplemental Table 5**).

**Figure 6:**
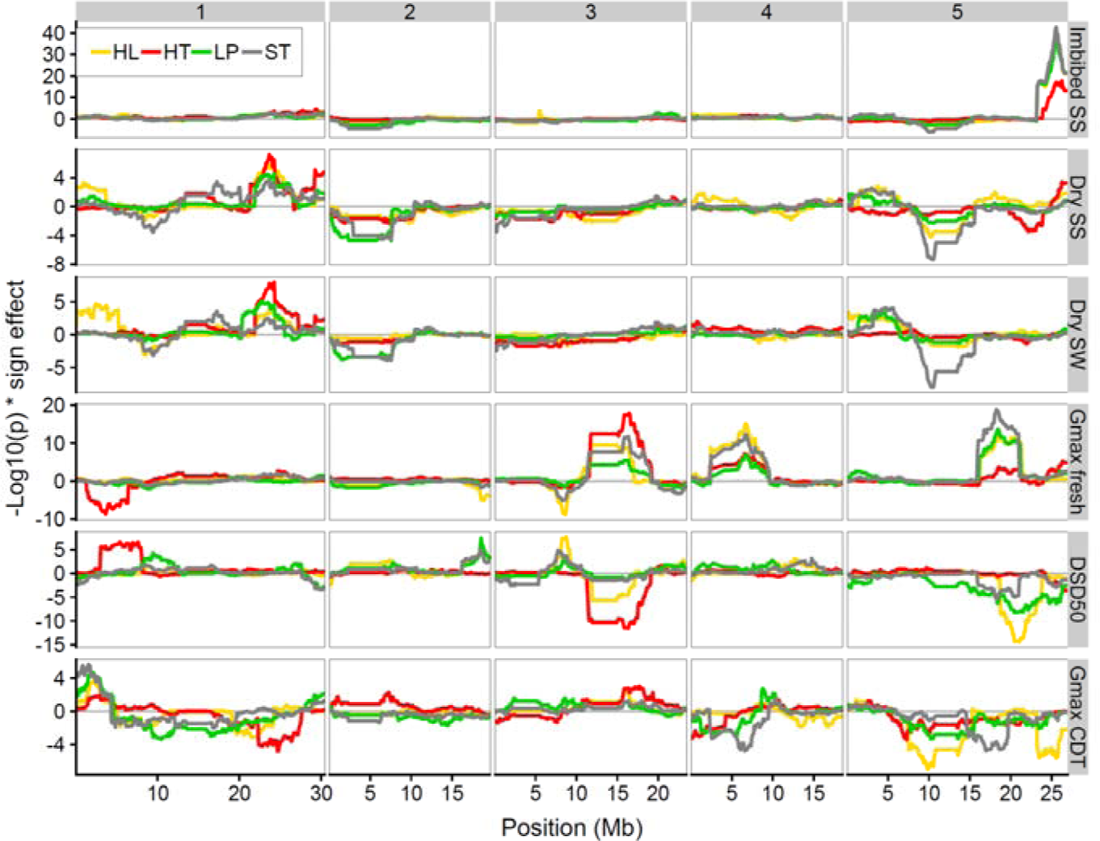
QTL profiles plot of seed and germination traits; Imbibed and dry seed size (SS), dry seed weight (SW), fraction of germination of fresh seeds (G_max_) and after controlled deterioration (G_max_ CDT), and days of dry storage required to reach 50% of germination (DSDS_50_). Chromosome numbers are show at the top. Genomic positions (x-axes) are indicated in mega base pairs (Mb) per chromosome. QTL effect direction and likelihood are indicated by significance (−log_10_) multiplied by the effect sign, on the y-axis (+ Bay-0 effect > Sha effect, - Sha effect > Bay-0 effect). The colors of the QTL distributions indicate the maternal environment; ST, grey; LP, green; HT, red; HL, yellow.

For imbibed seed size we detected one highly significant QTL at the right arm of chromosome 5, which was observed in all four ME datasets, suggesting that the explaining polymorphism underlying this QTL is a genetic determinant of seed size independent of maternal environment perturbation. For dry seed size several maternal environments QTLs were identified. Most QTLs were shared by at least two different ME’s, except some QTLs on chromosome 5, which appear specific to the HT maternal environment. Interestingly, dry seed weight showed a very similar QTL distribution pattern as dry seed size both in terms of location and effect. However, the HT-specific QTLs on chromosome 5 for dry seed size were not mirrored in the seed weight data set. In addition, the HL and LP ME QTL on the center of chromosome 5, that is shared with the ST ME conditions in the dry seed size dataset, is not observed in the dry seed weight QTL distribution.

For G_max_ of fresh seeds, QTLs were found on chromosomes 1, 3, 4, and 5. The QTL on chromosome 1 was only detected for the HT maternal environment, whereas contrarily the QTL found on chromosome 5 was almost absent in the HT maternal environment but present in the other ME’s. In the HT maternal environment, the DSDS_50_ QTL distribution revealed a QTL on chromosome 1 that co-locates with the G_max_ of fresh seeds QTL although it has an opposite effect. For the other maternal environments, no QTLs were detected at this genomic position. However, dry seed size and dry seed weight QTLs for HL, and G_max_ CDT QTLs for all ME’s were detected slightly upstream (left arm chromosome 1) and overlapped with the confidence interval of the G_max_ fresh QTL. Additional DSDS_50_ QTLs were detected on chromosome 3 for HT and HL and chromosome 5 for HL, LP and ST. For G_max_ CDT several diffuse small effect QTLs were observed lacking well-defined borders. Some of the QTLs co-located with QTLs found for seed size and seed weight.

Overall, we conclude that there is a complex genetic component that modulates the effect the maternal environment has on diverse progeny seed traits. Several of the QTLs were detected in multiple ME environments, which suggests that integrating diverse maternal environments into seed traits involves shared genetic loci (**Figure 6, Supplemental Table 5)**. Nevertheless, several QTLs were only detected for a single ME condition, suggesting that also ME-specific genetic components mediating seed trait values exist.

### QTL mapping of germination traits with QTL x E

Since the germination traits were measured in different germination environments (GE) using seeds derived from different maternal environments (ME), two QTL mapping approaches were used to explore QTL-by-environment interactions (QTL x E). We mapped QTLs for the seed germination traits AUC and t_50_ of seeds derived from mother plants that received ST, LP, HT, and HL environmental treatments. These seeds were exposed to salt, cold, heat, osmotic stress, or ABA during germination. Single trait multiple-environment linkage analysis was performed on the mean phenotypic values of the AUC across germination (GE) conditions, for each maternal environment (ME). As a second approach, all observations were fitted in a linear model, which allowed estimation of the interactions between the genomic marker, maternal environment, germination environment and stratification treatment.

The single trait multiple-environment mapping approach revealed many phenotypic QTLs distributed over the five chromosomes, for AUC and t_50_ (**Figure 7, 8, 9, 10, Table 1, Supplemental Tables 6 & 7**). For both AUC and t_50_, the co-locating QTLs were grouped in 13 main QTL clusters per trait (**Table 1**). The direction of the effects of the QTLs within these clusters was consistent across the environments and QTLs with both Bay-0 and Sha as high value allele were identified (**Figure 7, 8, 9, 10**). Yet, the distribution of allelic effects was much more equal between AUC traits, than for the t_50_ traits, with more positive allelic effects for the Bay-0 allele (**Figure 8, 10**). We observed variation in the number of QTLs, as well as variation in the explained variance of individual QTLs across the different ME’s, which is indicative for substantial QTL x E interaction. In addition, germination QTLs with environment specificity were identified. For example, for AUC, in the HT maternal environment data, several QTLs were identified on the left arm of chromosome 1 and chromosome 3, while QTLs at the bottom of chromosome 1 and top chromosome 5 are mostly specific to LP (**Figure 7**). For t_50_, the HT maternal environment showed the most specific QTLs in both position as well as strength/significance (**Figure 9, 10**). This shows that the genetic loci underlying trait variation can be modulated by the maternal environment and emerge as ME-specific QTLs. Of note, for t_50_ we observed a bias for higher trait values associated with Sha alleles and overall, the significance of t_50_ QTLs is less pronounced than observed for AUC (-log 10(p) range of ∼15 respectively ∼100).

**Figure 7:**
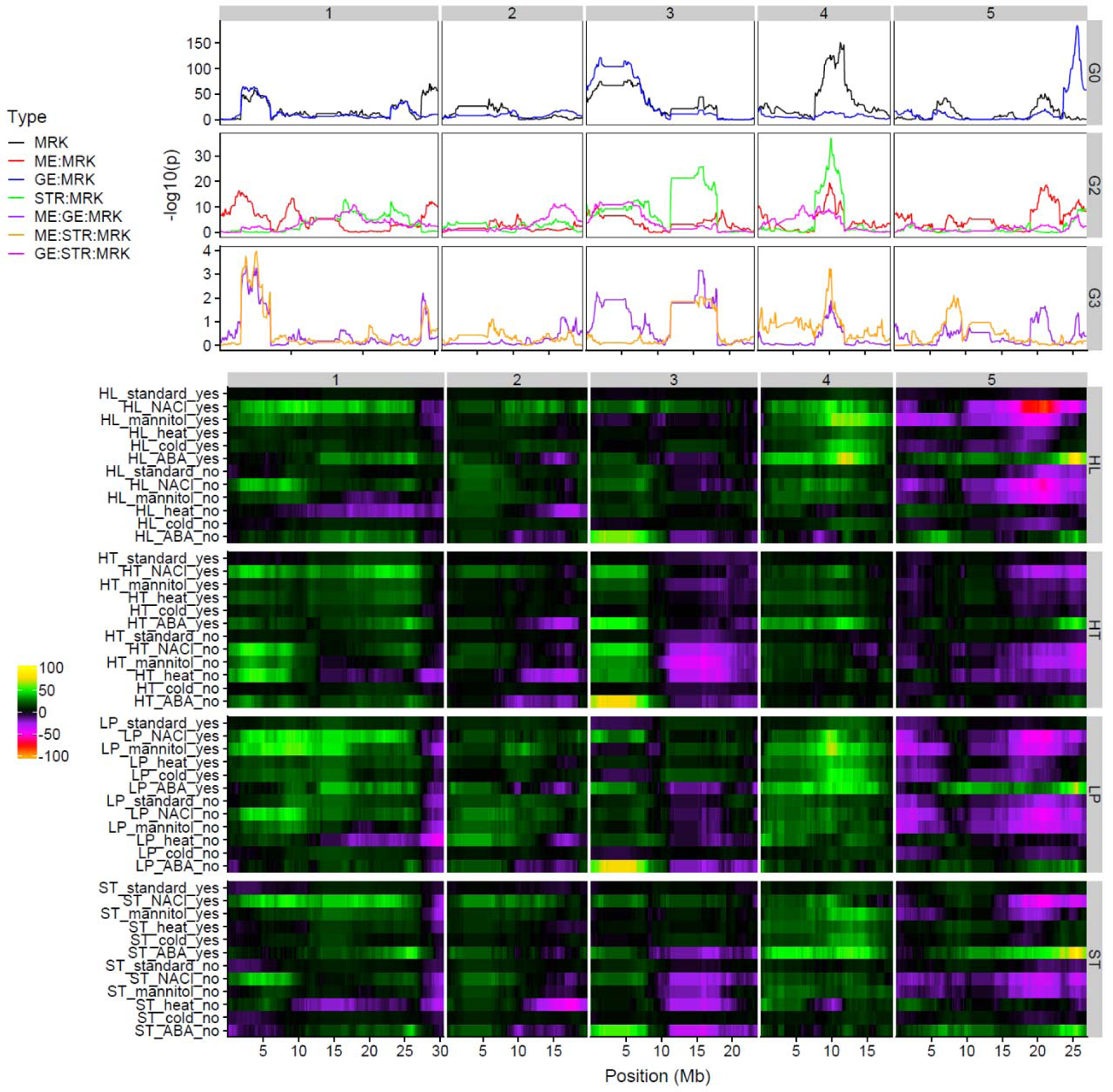
Distribution plot (A) and heatmap (B) of QTL profiles for AUC calculated by a single trait multiple environments linkage analysis approach. (A) The -log_10_(p-value) significance profiles for different components of the mixed model mapping approach are given, which are indicated by different colors: genotypic marker (MRK), maternal environment (ME), germination environment (GE), stratification (STR). Right-side label: G0 show the most significant effects: MRK and GE x MRK. Right-side label: G2 shows less significant effects: ME x MRK, STR x MRK and GE x STR x MRK. Right-side label: G3 show the least significant effects: ME x GE x MRK and ME x STR x MRK. (B) QTL profiles of the 12 germination conditions (Standard, NaCl, Mannitol, Heat, Cold and ABA, all with (yes) or without (no) stratification) are indicated in the rows per block of maternal environment conditions (standard, ST; low phosphate, LP; high temperature, HT; high light, HT). The direction and effect of the QTLs are indicated by the color scale; purple to orange (via red) indicates a higher trait value associated with the Sha allele (defined as negative effect) and green to yellow indicates a higher trait value associated with the Bay-0 allele (defined as positive effect). The white vertical lines delineate the chromosomes (A,B) Genomic positions (x-axes) are indicated in mega base pairs (Mb) per chromosome (chromosome numbers indicated on top of each block.

**Figure 8:**
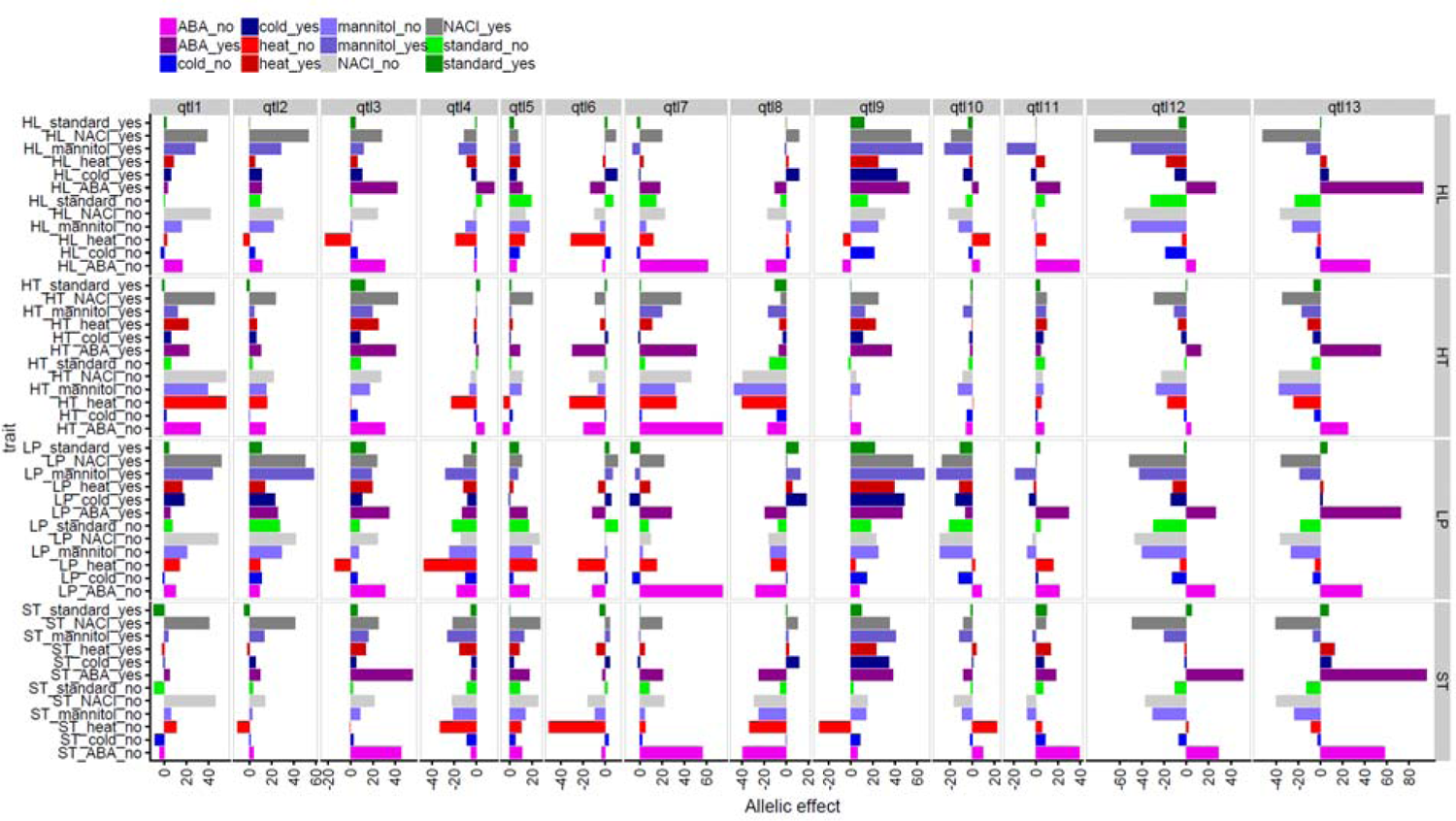
Allelic effects (AUC difference between the bay and sha alleles) of the 13 main identified QTL regions (indicated as qtl1 to qtl13 above the columns) detected for the AUC trait. X-axis shows allelic effect. Y-axis shows the AUC phenotype measured under different GE conditions (Standard, NaCl, Mannitol, Heat, Cold and ABA, all with (yes) or without (no) stratification) of seeds derived per ME (horizontal blocks; standard, ST; low phosphate, LP; high temperature, HT; high light, HT), also indicated by colors. Panel labels on the right indicate the ME conditions of the seeds. Positive signs indicate Bay-0 effect > Sha effect and negative signs indicate Sha effect > Bay-0 effect.

**Figure 9:**
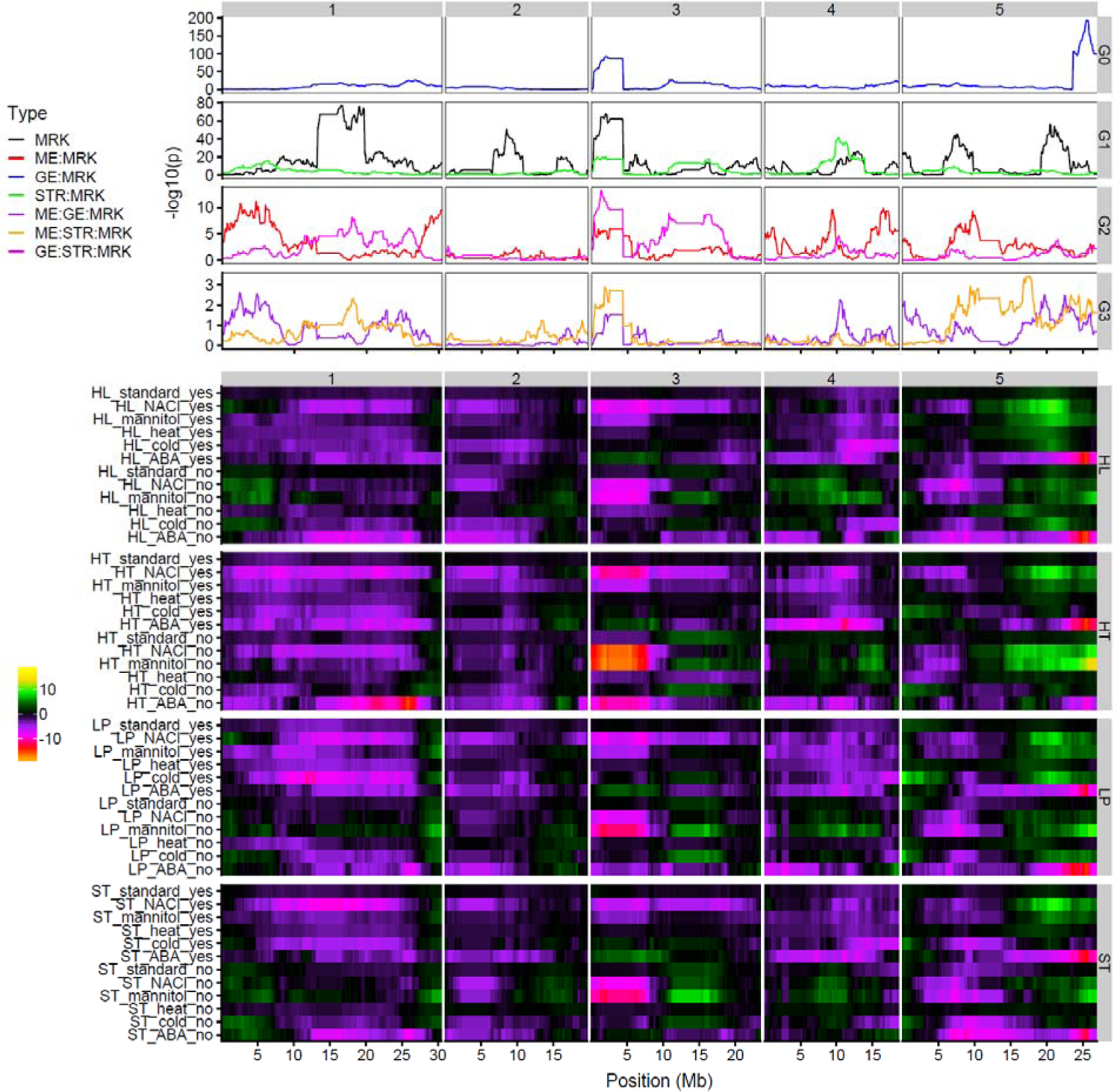
Distribution plot (A) and heatmap (B) of QTL profiles for t_50_ calculated by single trait multiple environments linkage analysis approach. (A) The -log_10_(p-value) significance profiles for different components of the mixed model mapping approach are given, which are indicated by different colors: genotypic marker (MRK), maternal environment (ME), germination environment (GE), stratification (STR). G0 show the most significant effect, GE x MRK. G1 the most significant after G0, MRK and STR x MRK. G2 shows less significant effects, ME x MRK, and GE x STR x MRK. G3 show the least significant effects, ME x GE x MRK and ME x STR x MRK (B) QTL profiles of the 12 germination (GE) conditions (Standard, Mannitol, NaCl, Heat, Cold and ABA, all with (yes) or without (no) stratification) are indicated in the rows per block of maternal environment conditions (standard, ST; low phosphate, LP; high temperature, HT; high light, HT). The direction and effect of the QTLs is indicated by the color scale; purple to orange (via red) indicates a higher trait value associated with the Sha allele (defined as negative effect) and green to yellow indicates a higher trait value associated with the Bay-0 allele (defined as positive effect). The white vertical lines delineate the chromosomes (A, B) Genomic positions (x-axes) are indicated in mega base pairs (Mb) per chromosome (chromosome numbers indicated on top of each block.

**Figure 10:**
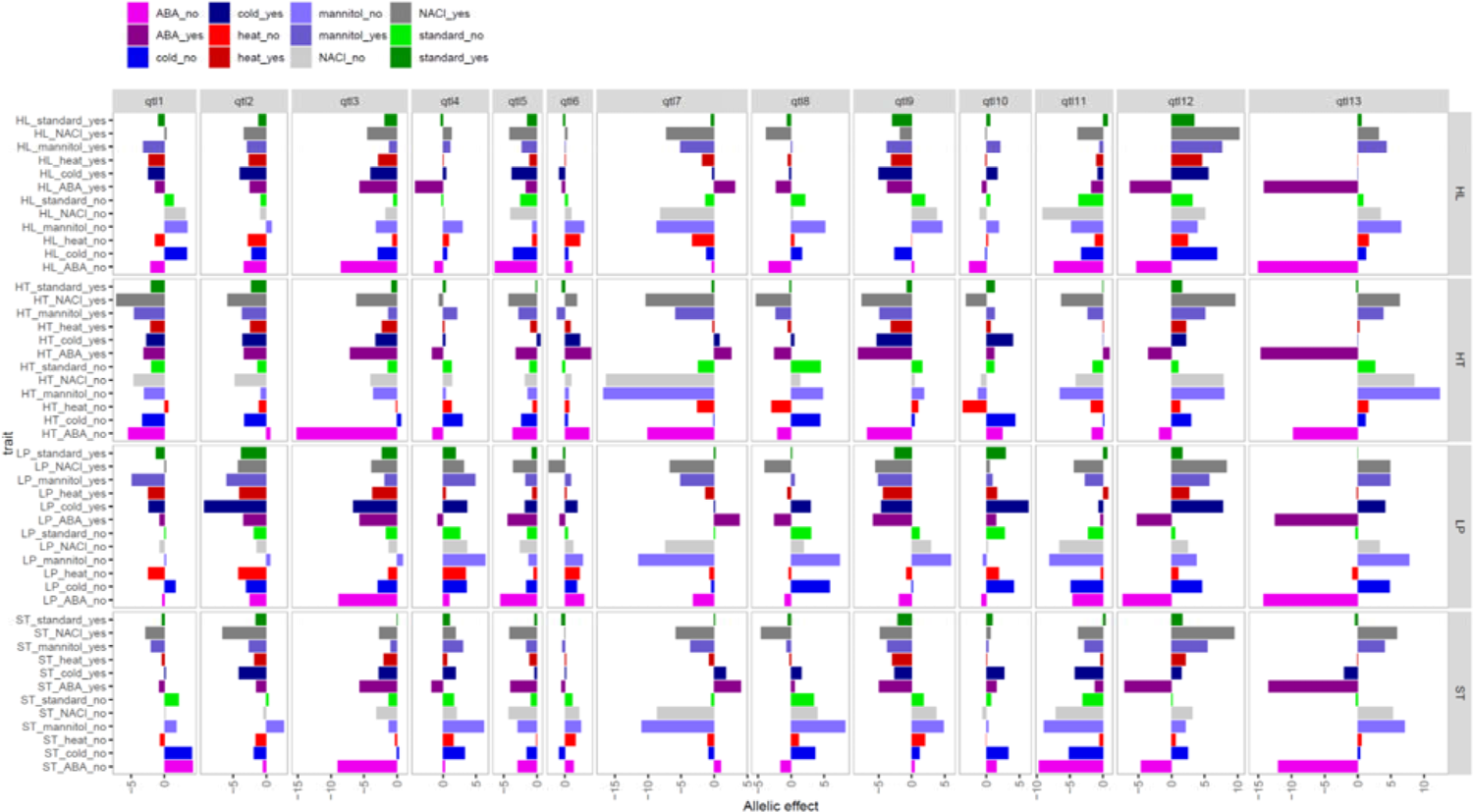
Allelic effects (t_50_ (hours) difference between the bay and sha alleles) of the 13 main identified QTL regions (indicated as qtl1 to qtl13 above the columns) detected for the t_50_ trait. X-axis shows allelic effect. Y-axis shows the t_50_ phenotype measured under different GE conditions (Standard, NaCl, Mannitol, Heat, Cold and ABA, all with (yes) or without (no) stratification) of seeds derived per ME (horizontal blocks; standard, ST; low phosphate, LP; high temperature, HT; high light, HT), also indicated by colors. Panel labels on the right indicate the ME conditions of the seeds. Positive signs indicate Bay-0 effect > Sha effect and negative signs indicate Sha effect > Bay-0 effect.

**Table 1:** Main QTL regions detected for the AUC and t_50_ phenotypes. QTL region name, marker ID (bin_marker) used for effect calculation, chromosome number, physical position (Mbp) and genetic position (cM) are given in columns.

|  | bin_marker | chromosome | physical.position.(Mbp) | genetic.position.(cM) |
| --- | --- | --- | --- | --- |
| qtl1 | RSM_1_3.25 | 1 | 3.25 | 13.59 |
| qtl2 | RSM_1_9.95 | 1 | 9.95 | 41.39 |
| qtl3 | RSM_1_26.05 | 1 | 26.05 | 94.68 |
| qtl4 | RSM_1_29.65 | 1 | 29.65 | 116.9 |
| qtl5 | RSM_2_6.75 | 2 | 6.75 | 20.24 |
| qtl6 | RSM_2_17.05 | 2 | 17.05 | 62.11 |
| qtl7 | RSM_3_3.35 | 3 | 3.35 | 10.44 |
| qtl8 | RSM_3_16.15 | 3 | 16.15 | 49.89 |
| qtl9 | RSM_4_10.35 | 4 | 10.35 | 52.51 |
| qtl10 | RSM_5_1.35 | 5 | 1.35 | 7.147655 |
| qtl11 | RSM_5_7.15 | 5 | 7.15 | 29.00167 |
| qtl12 | RSM_5_21.15 | 5 | 21.15 | 78.06281 |
| qtl13 | RSM_5_25.65 | 5 | 25.65 | 96.11812 |

## Discussion

It is well-known that the environment experienced by the mother plant during seed development and maturation affects progeny seed traits such as dormancy and germination [11]. The genotype-dependent response to environmental cues sparked studies on the interaction between maternal environment and genotype, and the effect on seed phenotypes [2, 18, 22, 34, 45, 49]. However, only a handful of studies took advantage of natural occurring genetic variation to identify the extent and effect of the maternal environment on seed traits at the genetic level and systematically assessed the G x E interactions. Postma and Agren [43] and, Kerdaffrec and Nordborg [44] identified changes in the effect size of QTLs associated to seed dormancy in Arabidopsis populations grown in field experiments. However, various factors such as photoperiod and temperature, change continuously, unpredictably, and simultaneously in the field. Therefore, experiments under controlled conditions, in which the effect of tweaking individual environmental cues without changing others, are useful to disentangle effects of individual environmental inputs precisely. In this context, we grew an Arabidopsis recombinant inbred line (RIL) population under four controlled conditions from flowering until seed harvest. Extensive phenotyping and QTL mapping of seed and germination traits of progeny produced under the different conditions revealed the effect of the maternal environment at the phenotypic and genetic level. In addition, we used fresh and after-ripened seeds, dry seeds, and imbibed seeds, stratified and non-stratified seeds and let these germinate under mildly stressful environmental conditions.

Overall, considerable phenotypic variation was observed between the parental lines as well as within the Bay-0 x Sha RIL population in response to both the germination environment and maternal environments (**Figure 1, 2**). In many instances, the phenotypes of the RILs extended beyond the parental values, indicative of segregating natural genetic variation in alleles that affect the trait values in a genetically additive manner. The effects of the maternal environment and germination environments on seed traits were in line with previous studies. For instance, temperature is a known key determinant in the timing and duration of key developmental phases including flowering and plant morphology, as well as reproductive development [50–52]. Warmer temperature leads to earlier flowering and subsequently earlier seed set [9, 51, 53]. In line with other studies[12, 15], the high temperature maternal environment promoted germination by reducing dormancy levels (**Figure 1**). Altogether, our complex multi-variate approach demonstrates that multi-environment analysis is a valuable tool for understanding the genetic control of seed and germination traits, and other traits alike.

Larger seed size and seed weight was observed under HL (**Figure 1**), in line with findings in other genetic backgrounds [2, 17]. In several studies, seed size has been associated with seedling vigor and faster germination [14]. However, the low correlation between seed size and seed performance found in this study (**Figure 3**) shows that this relation is at least complex. This is in accordance with the poor correlation between tomato seed traits and seedling performance, in different maternal environments reported by Geshnizjani et al. (2020). Seeds that matured under HL conditions did show a reduced percentage of germination when exposed to NaCl, Mannitol or heat (**Figure 2)**. This could be explained by either a higher water requirement of these seeds to complete germination compared to smaller seeds, for which water absorption can occur faster, or by a disrupted osmotic balance which may complicate water uptake by the seeds. These findings are in line with an earlier observation that an increase in resources in the maternal environment, caused by high light, decreases the fraction of seeds with early germination [54]. When the mother plant experiences non-optimal conditions, such as low phosphate, offspring can become more dormant [2, 9]. The low phosphate environment also affected seed traits, such as germination in mannitol (**Figure 2**), which resulted in LP-sensitive QTLs at the top of chromosome 5 (**Figure 7, 8, 9, 10**). Under phosphorus starvation conditions, the allocation of phosphorus might occur at the expense of its storage in seeds, although studies have reported that mainly yield, rather than seed quality, is affected under resource limiting environments [19]. Together, these results indicate that suboptimal environmental conditions experienced by the mother plant, that are transmitted to the progeny, can proposedly be translated into different functional strategies, such as promotion or (temporary) inhibition of seed germination, allowing the seeds to adjust their germination behaviour to fit their immediate environment [55].

Besides the dominating effect of the seed’s germination environment on the various germination traits (**Figure 2**), the maternal environmental conditions also affected germination traits in the RILs, indicated by genotype-by-maternal environment (G x ME) interactions (**Figure 4, Table 1**). Such interactions were also reported in other studies [2, 18, 44] and are supported by this study through the analysis of variance, which estimated that, overall, approximately 40% of the seed germination variation in the RILs was due to G x ME (**Table 1**). Together, these results show that there is a genetic contribution, and natural variation therein, to the effect that the maternal environment has on the expression of progeny seed quality and germination traits. Our quantitative genetic study identified many seed quality and germination QTLs across the different ME’s and GE’s. We identified 13 QTL clusters (**Figure 8, 10, Table 1**), which correspond to genomic locations of previously identified QTLs controlling germination under standard and stress conditions [8, 56]. The effect of the maternal environments was mostly observed as change in allelic effect size of the detected QTLs, which is the most common type of interaction for QTL x E [33]. For marker-assisted selection in breeding approaches, such QTLs present the advantage of enhancing germination under various conditions. For these QTLs, alleles promoting germination can be queried from parents having opposite phenotypes.

The maternal and germination conditions significantly attributed to the magnitude-of-change of the QTL effects (**Figure 7, 8, 9, 10**). Common germination QTLs across maternal environments showed higher significance scores than ME specific QTLs. However, some ME specific QTLs could be detected, for example in the HT ME, yet some could easily be missed due to not reaching the significance threshold under these conditions. The result of the QTL analyses, and the extensive QTL x E interactions identified, illustrate the dynamics of the contribution of the genetic architecture to responses to different maternal environments. The fact that both common and maternal environment-specific QTLs were identified, suggests that studies that assess multiple environments like here presented could be more often considered to fill gaps in our understanding of gene regulatory networks, in response to environmental perturbations.

Taken together, our complex multi-variate approach demonstrates that multi-environment analysis is a valuable tool for understanding the genetic control of seed and germination traits, and other traits alike. We showed that the maternal environment, and the environmental condition during germination, prominently affect seed traits and seed germination, and that extensive natural genetic variation exists underlying the modulation of these effects. The extent of QTL x E observed because of genetic interaction with the maternal and germination environments strengthens the need for contrasting multi-environment studies to reveal the genetic mechanisms underlying phenotypic plasticity. Such studies are particularly relevant in the context of climate change [57, 58], to understand the fundamental mechanisms of plant adaptation and acclimation to environmental change [20], and also for breeding purposes, to safeguard future crop production in changing environments.

## Material and methods

### Plant materials and growth conditions

In total 165 lines of a Arabidopsis Bayreuth (Bay-0) x Shahdara (Sha) accessions recombinant inbred line core collection [46, 47] were used, including the parental lines. All lines were sown on imbibed filter paper, placed in individual petri dishes and stratified (4 days at 4°C in the dark). The sowing of the seeds differed in time based on the estimated flowering time of the RILs observed in previous experiments [8] to synchronize flowering. The seeds were then allowed to germinate in climate-controlled incubators with continuous light at 22°C. At the time of radicle protrusion, 16 seedlings per line were transferred to Rockwool blocks, with one seedling per block. The plants were further grown in a controlled climate room under long day conditions (light/darkness cycle of 16h/8h) at 22°C/18°C with a light intensity of 150 μmol m^-2^ s^-1^ and 70% relative humidity [2]. The plants were automatically watered with standard nutrient solution (**Supplemental Table S1**) three times a week.

When plants flowered, the stems and branches were cut in order to ensure development of all seeds occurred under the defined controlled mild stress condition (i.e. maternal environment; ME). Three to four plants per line were transferred to different climate cells set under different controlled environmental stresses being: low phosphate nutritive solution (LP; 0.01 mM), high temperature (HT; 25°C/23°C) and high light (HL; 300 μmol m^-2^ s^-1^), or were maintained in the standard growth conditions (ST; 20°C/18°C, 150 μmol m^-2^ s^-1^, 0.5 mM phosphate) [2]. The plants were kept in these conditions until seed harvest. When all plants produced a sufficient amount of fully matured seeds, seeds were bulk-harvested from 3-4 plants per genotype. A fraction of the fresh harvested seeds was dried and stored at −80°C in sealed 2 ml tubes, while the remaining seeds were stored in paper bags placed in a cupboard at ambient room temperature to allow after-ripening.

### Phenotyping

Seed phenotyping was performed as described previously [2]. Seed size was determined by taking pictures of approximately 500 seeds on white filter paper using a Nikon D80 camera. Pictures were analyzed using ImageJ. The same seeds were transferred into weighing cup and weighed with an AD-4 Autobalance (PerkinElmer, Inc.). Single seed weight was subsequently determined by dividing the total weight by the number of seeds.

Germination experiments were performed using the GERMINATOR set-up described in [59]. Seeds were sown on two layers of blue filter paper (Anchor paper company, St Paul, MN, USA; www.seedpaper.com) with 48 ml of demi-water. Up to 6 seed batches were sown on parts of the same filter paper. Automatic scoring of seed germination was performed using a mounted camera system. Pictures were taken one to three times a day, for 5 up to 10 days after sowing, until green cotyledons became visible.

The curve fitting module of the GERMINATOR setup was used to analyze the general cumulative germination data. G_max_ was measured as the total seed germination percentage at each time point. T_10_ and t_50_ values were calculated as the time needed to reach 10% respectively 50% of germination of viable seeds. The calculation of the area under the germination curve (AUC) was extended to 300 hours to capture the full phenotypic variation under all germination conditions. The germination tests were performed using approximately 50 seeds per experiment. Two independent experiments were performed to obtain replicated phenotypic values.

The germination potential of the freshly harvested seeds (G_max_ fresh) and release of primary seed dormancy were determined by performing weekly germination experiments. Seeds were considered fully after-ripened if the percentage of germination reached more than 90% in two consecutive germination experiments. Fully after-ripened seeds were transferred to sealed Eppendorf tubes and stored at −80°C to prevent loss of viability during storage. The DSDS_50_ was calculated as the number of days of seed dry storage required to reach 50% germination [24].

The vigor of the seeds was assessed using fully after-ripened seeds by germinating the seeds in twelve different germination conditions. These germination experiments were initiated when more than 80% of the lines were fully after-ripened, as described above. Seeds were germinated using demineralized water in standard germination conditions. Germination experiments in sub-optimal conditions were conducted at high (32°C) and low (10°C) temperatures, under osmotic stress (−0.6 MPa mannitol; Sigma Aldrich); under salt stress (100 mM NaCl; Sigma Aldrich; ∼-0.45 MPa) and in presence of ABA (0.25 µM ABA, Duchefa Biochemie). Germination experiments in these conditions were performed with and without stratification. Stratification consisted of storage of the sown imbibed seeds in the dark for 4 days at 4°C prior to transfer to the light (i.e. allowing germination). Since stratification can induce variation in stress sensitivity, we adjusted the concentrations for the NaCl and ABA treatments to 125 mM NaCl and 0.5 µM ABA for the experiments with stratification. NaCl and mannitol stress and ABA treatments were performed by adding solutions of the indicated concentrations to the filter paper instead of solely demi-water prior to stratification.

Seed longevity was assessed by a controlled seed deterioration test. To this aim, dry seeds were incubated at 40°C at 85% relative humidity in a closed tank in the presence of a saturated ZnSO4 solution. After five days, seeds were removed and germinated in standard conditions as described above.

### Data analysis

Previous studies showed no difference in the QTL mapping when performed using either transformed or non-transformed germination data [8]. Therefore, due to the large number of traits, all analyses were performed on untransformed data, except for DSD_50_ which was square root transformed. Figures were made using the R [60] package GGplot2 [61]. Heatmaps were made using the R package gplots [62]. ANOVA analysis was performed for phenotypic mean comparisons of the seed traits measured in the RILs grown under the four maternal environments. Post-hoc Tukey test was then used at the confidence level of 0.99 to determine pairwise group significant differences.

### Heritability

For each trait in each maternal environment (ME), the broad-sense heritability (*H2*) was calculated from estimated variances: *H2 =*σ *^2^G / (*σ *^2^G+*σ *^2^E)*

Where *σ2G* is the genetic variance and σ*2E* is the environmental variance. The variance component analysis was analysed using a two-step mixed model approach (REML) from the preliminary single environment analysis in Genstat (18th Edition). Genotype and replicate were set as random effects in the model.

### QTL analysis

QTL mapping was performed using a genetic map for the Bay-0 x Sha population derived from RNA-seq data [47]. Briefly, the genotyping of 160 RILs resulted in the identification of 1059 polymorphic markers between the two parental lines. A general linear multi-marker model was used with a forward mapping approach, with a maximum of 9 cofactors and 40 marker window size (∼4Mbp, whole window, ∼2Mbp to each side). All QTL profiles can be found in AraQTL [48].

## Acknowledgements

This work was supported by Technology Foundation (STW), which is part of the Netherlands Organization for Scientific Research (NWO) (EARS, HN, LAJW, JAR, WL).

## Author contributions

HWMH and WL conceived the study. EARS, LAJW and JAR performed the experiments. BLS, EARS and HN analyzed the data. BLS, EARS, MvZ and WL wrote the manuscript with input from all co-authors.

## Supplemental data

**Supplemental Table S1:** Information about measured traits and plant growth conditions.

**Supplemental Table S2:** List of all measured traits

**Supplemental Table S3:** Average phenotypic values for all seed and germination traits measured in the parents and recombinant inbred lines grown under standard, low phosphate, high temperature and high light.

**Supplemental Table S4:** Correlation between all traits

**Supplemental Table S5:** QTL profiles for seed traits

**Supplemental Table S6:** QTL profiles for AUC

**Supplemental Table S7:** QTL profiles for t_50_

## Supplemental Figures

**Supplemental figure 1:**
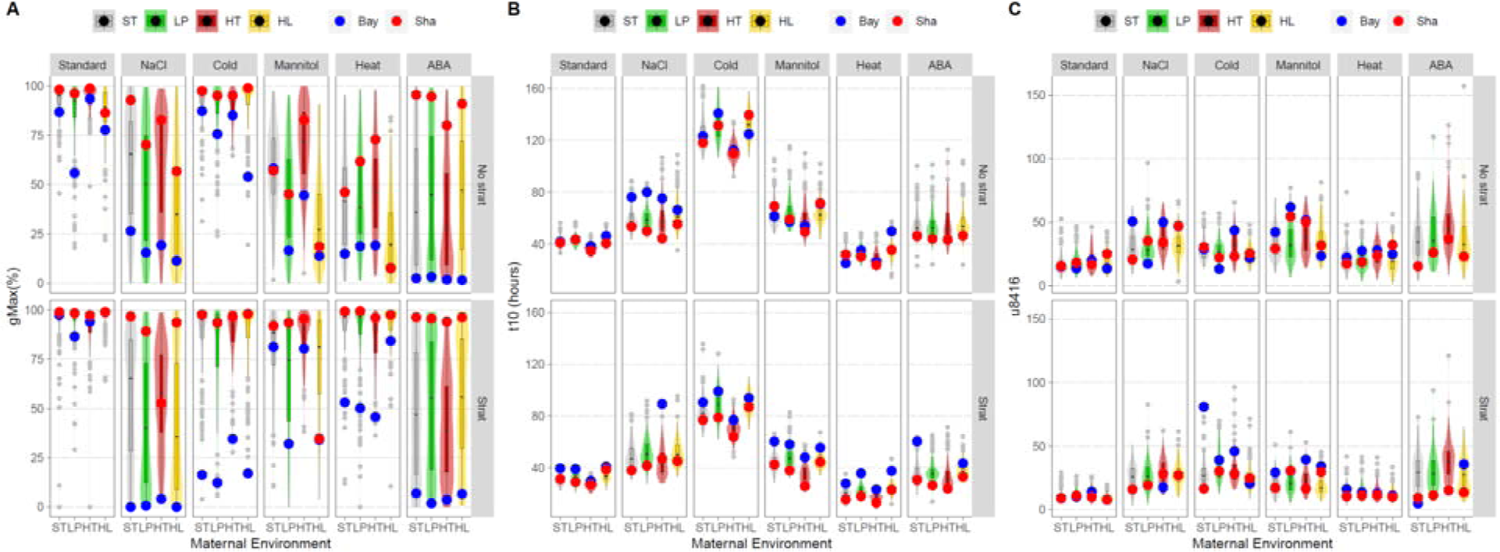
Effect of the seed maternal and germination environment on seed vigor traits of RIL population genotypes and parental lines. (A) G_max_, (B) t_10_ values and (C) u_8416_. The phenotypic values of the parental lines, Bay-0 and Sha are indicated with blue and red dots, respectively. G_max_, t_10_ and u_8416_ were assessed using fully after-ripened non-stratified seeds (upper row) and fully after-ripened stratified seeds (lower row) and were exposed to salt, cold, heat or osmotic stress or ABA. Boxes indicate boundaries of the second and third quartiles of the RIL distribution data. Black bars indicate median and whiskers Q1 and Q4 values within 1.5 times the interquartile range. Outliers are represented by grey dots. Violin plots designate phenotypic distributions. Colored shadings represent the different maternal environments; standard (control) conditions (ST, grey), low phosphorus (LP, green), high temperature (HT, red) and high light (HL, yellow).

**Supplemental figure 2:**
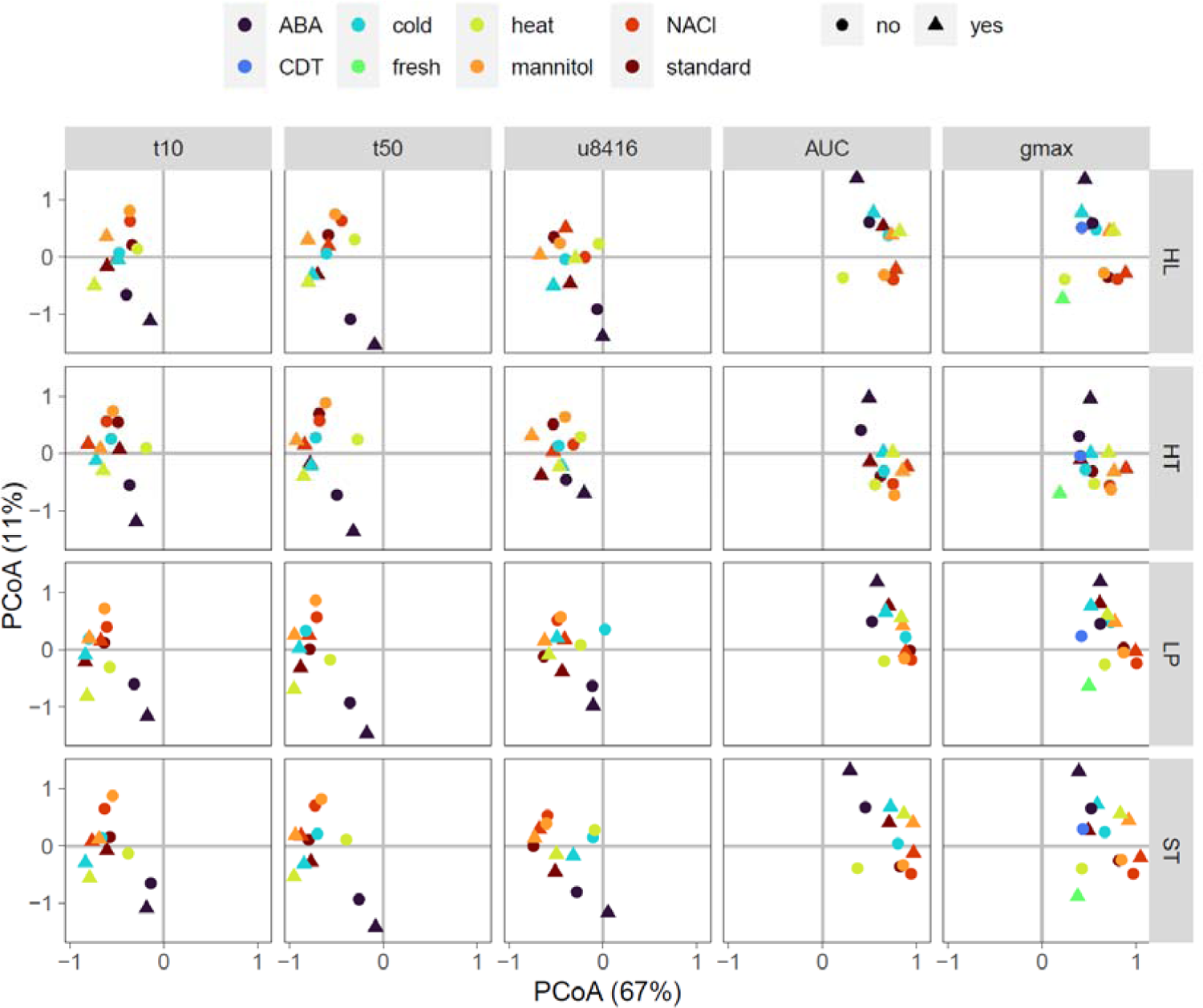
PCoA of the phenotypic values of the germination traits. (RIL and parental) between the maternal environment and germination environments. PC1 is shown on the x-axis, including explained variation, PC2 is shown on the y-axis, including explained variation. Germination environments indicated by color, stratification (yes/no) indicated by shape, trait indicated by the labels on top of the panels, maternal environment indicated by labels on the right of the panels.

