## Supplementary figures and images for "Modulation of quantitative trait loci for *Arabidopsis thaliana* seed performance by the maternal and germination environment"

### Supplemental figure 1

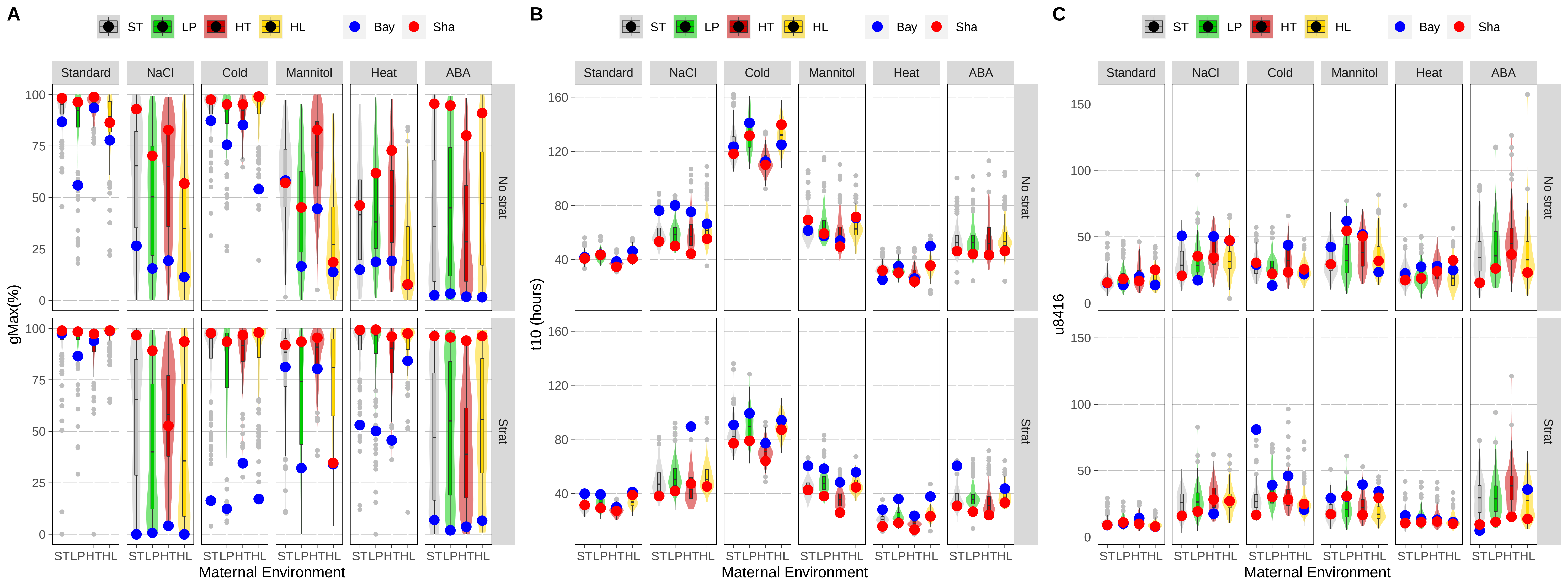

### Supplemental figure 2

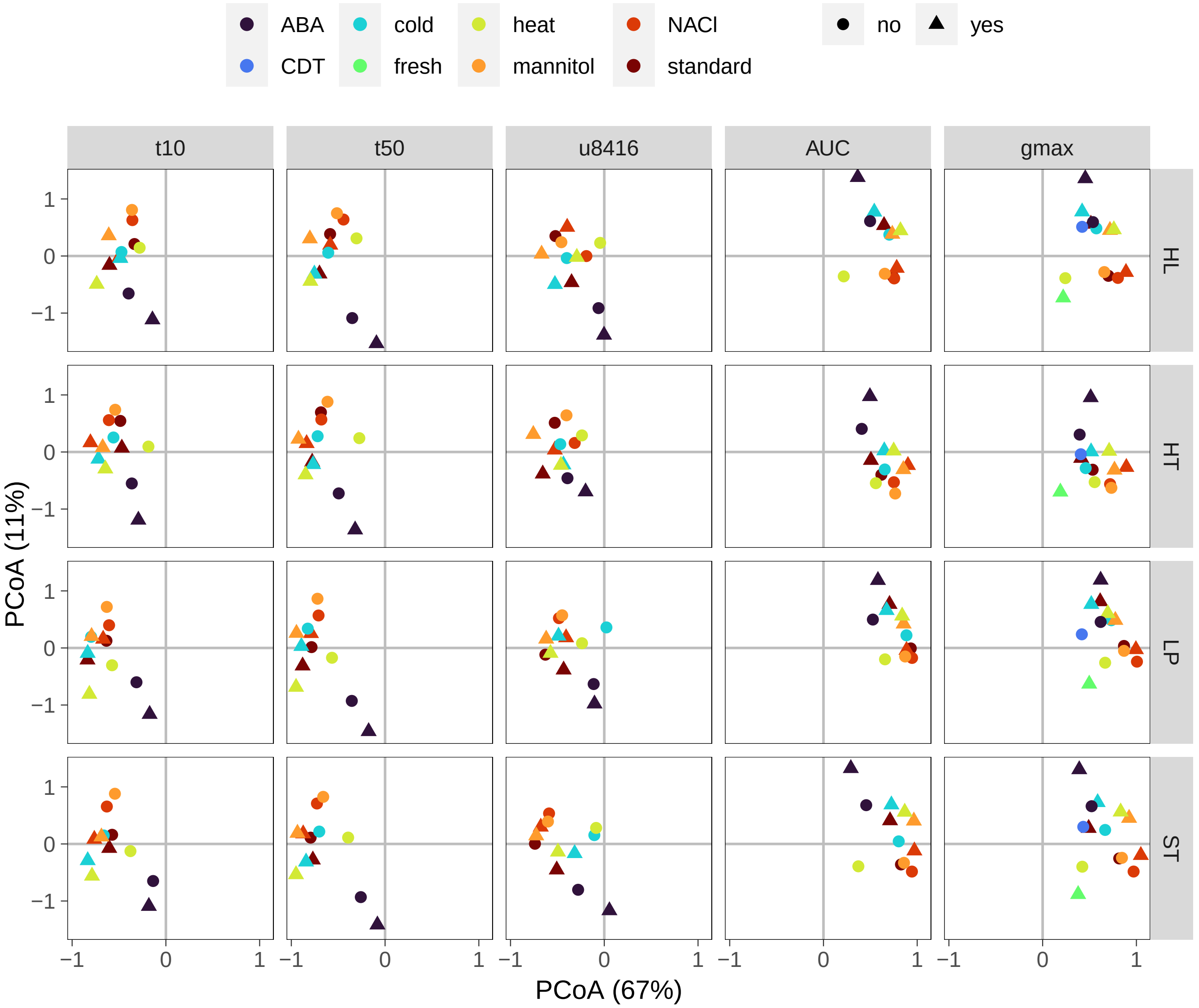
